# MaD: Macromolecular Descriptors for Integrative Dynamic Modeling Supported by Cryo-EM Data

**DOI:** 10.1101/2022.06.22.497181

**Authors:** Sylvain Träger, Matteo Dal Peraro

## Abstract

The determination of atomic structures of large flexible systems remains a challenging task despite the recent advances in cryo-electron microscopy (cryo-EM) and *de novo* protein structure prediction. Few hybrid methods truly consider dynamics because deriving appropriate conformational ensembles and scoring functions robust to conformational differences is not trivial. We present here Macromolecular Descriptors (MaD) which, inspired by traditional computer vision concepts, remedies some of these limitations. MaD identifies feature points and encodes local structural information around them into resolution- and conformation-invariant descriptors. Efficient matching of descriptors from cryo-EM densities at low and medium resolution with those of high-resolution component structures yields a robust and accurate assembly prediction that does not require other experimental or computational input. Fast, scalable and easy to use, this method is able to incorporate native dynamic features as extracted from molecular simulations and identify the models that best match the target electron density. Therefore, MaD addresses some of the unanswered needs of the community in terms of the integrative modeling of large and flexible macromolecular complexes.

## Introduction

Structural biology is at the frontline of scientific research thanks to its ability to elucidate cellular functions from structural knowledge of the underlying macromolecular assemblies. Typically made up of several protein components, they exhibit complex dynamics in order to fulfil their task in their natural cellular environment. Traditional structural biology techniques such as X-ray crystallography, nuclear magnetic resonance (NMR) and more recently cryo-electron microscopy (cryo-EM) and tomography (cryo-ET) are typically employed to obtain discrete structural snapshots but still struggle as the molecular weight and flexibility of the structure increase. Of note, the average resolution achieved through single particle cryo-EM still has not breached the atomic barrier (5.28 Å in 2021^1^), while subtomogram averaging has improved significantly, now achieving subnanometer resolutions^2,3^. Atomic structures from either techniques are therefore not guaranteed in general and the approach to model structures from their data subsequently depends on their resolution^4^. When atomic resolution is not achieved, the modeling burden is transferred to computational, integrative methods. High resolution structures of individual components, often obtained from X-ray crystallography but now also available from computational predictions^5,6^, and a medium/low resolution cryo-EM/ET density map of the assembly are fed to a multi-resolution integrator that aims to minimize a predetermined scoring function, whose global minimum theoretically corresponds to high-confidence models of the assembly. If available, spatial information from chemical cross-linking or mutagenesis experiments may be considered to limit uncertainty and facilitate convergence^7,8^.

However, existing approaches mostly adopt a rigid perspective of the target assemblies, postponing any consideration of dynamics to local refinement procedures^4,9,10^ after the prediction^11^. This is in striking contrast with the observation that components may behave differently in isolation than in their oligomerised form^11–14^. Despite the acceptance that molecular structures exist as a continuum of states^15^, the practical realisation of this idea is still lacking. Cryo-EM still mostly results in a series of discrete structural snapshots. Recent methods are able to derive continuous conformational changes at the single particle level in some cases^16^, but related 3D reconstructions are usually limited to subnanometer resolutions at best. As a result, the modeling community is in dire need of tools able to consistently integrate dynamic information to guide modeling in general and reach an exhaustive, atomistic overview of flexible systems.

Early integrative methods aiming to predict assemblies on the basis of cryo-EM data revolved around summarizing input structures to a set of features, turning the prediction task into a point-matching problem^17–19^. Due to the difficulty of detecting reliable features in experimental cryo-EM data, more recent methods combine a reduced number of features with different optimizers^20–22^, or instead represent structures as Gaussian mixture models^23,24^. Other methods opted for different schemes to scour the complex search space including a degree of flexibility^25^ or discretised space explored with Fast Fourier transforms^26,27^. In general, the nature of scoring functions varies, but generally include terms related to shape complementarity, structural clashes or protrusion out of experimental densities. These terms are incompatible with conformational differences between input structures, which combined with the inherent noise and errors within experimental cryo-EM/ET data result in a painstaking modeling process. As a result, many researchers still resort to manual and visual predictions, often aided by the local optimization tool Fit-In-Map from Chimera^28^. A previous method from our group is able to incorporate explicit subunit dynamics during prediction^12–14^, but remained limited to symmetrical homomultimers to avoid the excessive computational complexity of heteromultimeric assemblies.

We introduce here *Macromolecular Descriptors* (MaD), a method that revisits feature-based approaches by taking inspiration from local feature descriptors from computer vision. Descriptors are routinely applied in a variety of tasks, ranging from image registration, reconstruction or object detection^29,30^, with some descriptors built specifically for biomedical images^31–33^. In general, descriptors are built from local regions surrounding feature points, processed for rotation invariance, high local specificity and robustness towards noise or various transformations. We adapted concepts inspired by the well-performing scale-invariant feature transform (SIFT) descriptors^34–36^ towards integrative modeling tasks on the sole basis of cryo-EM data and molecular dynamics (MD) simulations, enabling the dynamic prediction of large assemblies efficiently. We first showcase the steps of assembly prediction before we demonstrate MaD’s predictive abilities over many experimental examples from cryo-EM/ET at various resolutions. We compare our approach to existing methods and demonstrate its robustness to conformational mismatch between input structures. Combined with previous tools we developed and with classical MD simulations, we further show how descriptors scale to conformational ensembles and effortlessly identify compatible structures.

## Results

### Assembly prediction using Macromolecular Descriptors

In the typical application, descriptors are generated for a cryo-EM map of a macromolecular complex at non-atomic resolution (> 5 Å) and high-resolution structures of its components. Efficient descriptor matching then lead to the identification of translation and rotation operations enabling the localization of components within the experimental density map, yielding atomistic models of the complex (see Methods section). First, component structures are converted to density maps matching the resolution of the target assembly, similar to previous methods^27,37^. Anchor detection (**Figure 1a**) consists in identifying a set of stable anchor points, which correspond to voxels with information-rich surroundings where large changes in intensity are present, enabling their reliable detection even in presence of noise or transformations. In practice, an anchor is a local maximum in a density grid after filtering with a Laplacian of Gaussian (LoG). Of note, the LoG filter has been used previously in hybrid modeling software to enhance shape contrast^38^. The second step of orientation assignment (**Figure 1b**) focuses on building spherical histograms from gradient vectors within a spherical patch around anchors, following the zonal equal sphere partition scheme^39^ (**Figure 1b**, grey spheres). Rotation matrices *R*_*c*_ and *R*_*a*_ are obtained for the component’s and assembly’s anchors, respectively, by placing the bins with the highest vector counts at key locations on the histograms (**Figure 1b**, red and orange arrows). This step sets anchors in a rotation invariant referential system and enables their comparison (**Figure 1b**, center). At this point, all the information needed to place components within the density of the target assembly is now available. The component may be translated using a vector *T*_*c*→*a*_ connecting one of its anchor to one of the assembly, rotated to a rotation-invariant reference frame by applying *R*_*c*_ and finally into the referential of the cryoEM data by applying the inverse of *R*_*a*_ (**Figure 1c**), granted that the chosen anchor pair matches in location and orientation.

**Figure 1:**
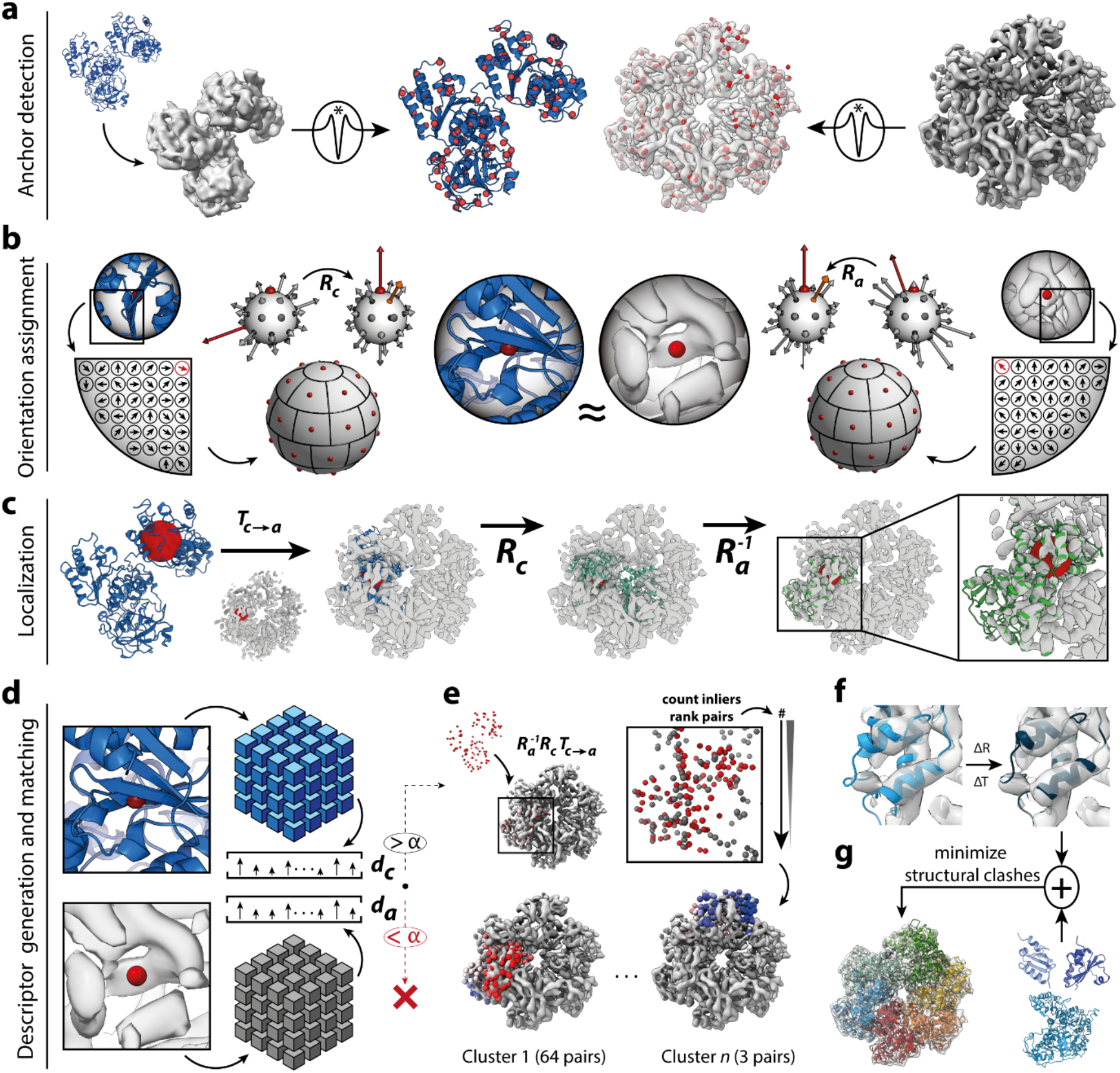
Integrative modeling with MaD (Macromolecular Descriptors). **a**, High-resolution structures are converted to simulated density maps at the same resolution of the cryo-EM data. Anchors are detected as local maxima after applying a Laplacian of Gaussian filter. **b**, A spherical patch is extracted around each anchor and a spherical histogram is built from the gradient vectors computed inside the patch. A rotation matrix is derived by rotating the two bins with highest vector counts at key locations on the histogram (red and orange arrows). Anchors are now rotation invariant and comparable. **c**, A component is localized in the cryo-EM density map by applying a translation vector *T*_*c*→*a*_ between anchor coordinates, a matrix *R*_*c*_ from orienting the component’s anchor and the inverse of *R*_*a*_ to bring the component in the referential of the cryo-EM data. **d**, To identify matching anchor pairs, descriptors are generated by subdividing the patch into 64 subregions. Spherical histograms are computed in each subregions before being flattened into a unidimensional vector. Descriptor pairs with a cross-correlation above a confidence threshold are selected. **e**, Component anchors are transformed into the referential of the cryo-EM map for each selected pair, which are then ranked according to their correspondences. Pairs are then clustered according to the final localization in the cryo-EM map. Solutions are extracted from the most populated clusters. **f**, Solutions are refined along density gradients. **g**, By adding the solutions of other components, assembly models are built by minimizing structural clashes and combining solutions.

The identification of such pairs is enabled by constructing descriptors for each anchor (**Figure 1d**). A cubic patch of the same diameter as in the previous step is extracted after orientation assignment and divided into an 4×4×4 array of subvolumes. This takes inspiration from the receptive fields of complex neurons within the primary visual cortex, which allows for gradient information to shift over small areas^35^ while enforcing information to be at a specific location around the anchor. As such, this process provides discriminative power to descriptors as well as robustness to noise and transformations such as conformational differences between input structures. Spherical histograms are built in each subvolume and the final descriptor is obtained by converting them to a unidimensional vector. Identification of matching pairs is achieved through cross-correlation, where pairs above a confidence threshold are selected.

Descriptors only consider a small structural patch and thus constitute a local measure. To score candidates at the level of the whole structure, we proceed to a global matching step (**Figure 1e**). Component anchors are transformed into the cryo-EM map according to the *T*_*c*→*a*_, *R*_*c*_ and *R*_*a*_ associated to each selected descriptor pair. The repeatability *R*, or the number of inliers between the anchors of the component and those of the assembly is computed and pairs are ranked accordingly. A fast hierarchical clustering step ensues, where pairs are allowed to vote for discrete localizations in the cryo-EM map obtained independently. A list of candidate solutions is extracted from the most populated clusters and refined using a procedure similar to Chimera’s Fit-In-Map tool^28^ (**Figure 1f**) to fix inaccuracies from anchor coordinates and spherical histograms. A solution *i* is finally scored as follows:

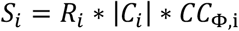

where *R*_*i*_ is the repeatability or percentage of corresponding anchors, |*C*_*i*_ | is the size of its cluster, and *CC*_Φ,i_ a global cross-correlation with the cryo-EM map. Our scoring function thus serves as a multi-level similarity measure. The procedure is typically repeated for other components or members of a structural ensemble. Finally and in general, the discrete nature of solutions and the limited amount of false positives enables a fast assembly building process achieved by a simple combinatorial minimization of structural clashes (**Figure 1g**).

### Application to homo- and hetero-multimeric assemblies at various resolutions

We first applied our method to a range of experimental assemblies at various resolutions and spanning different multimeric states. First, we modelled the NMDA receptor from an initial cryo-EM map at a resolution of 6 Å. This asymmetric heteromultimer is made of two GluN1, one GluN2A, one GluN2B subunits and a Fab domain bound to the latter. MaD reconstructed the assembly by docking and scoring each component independently. Of note, MaD was able to find both locations of GluN1 from either copy without false positives (**Figure 2a**, blue and green). GluN2A and GluN2B are correctly located with GluN2B being alternatively localized at the site of GluN2A. The Fab domain is easily placed as well despite the experimental density displaying poor quality in that region, leading to the identification of two false positives (**Figure 2a**, bottom). In this case, the MaD score enables a better discrimination of false positives compared to the traditional cross-correlation. Model building yields few models, with the first and best model having a Cα-RMSD of 1.09 Å with the reference structure and a cross-correlation of 0.90 with the experimental density (**Figure 2a**, right).

**Figure 2:**
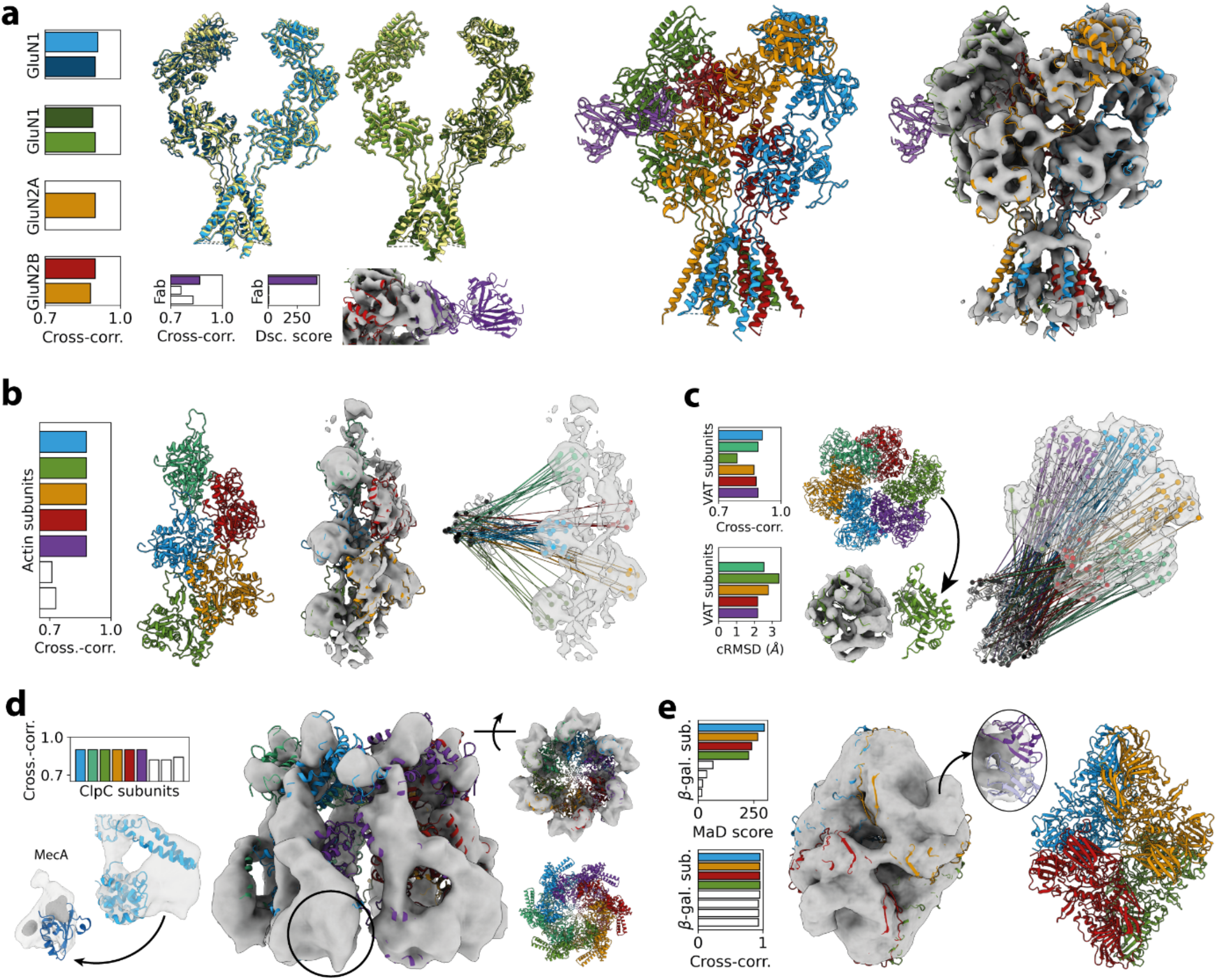
Applications to selected assemblies at various resolutions. **a**, The NMDA receptor (EMD-8584 at 6 Å, PDB 5UP2) consisting of two GluN1, one GluN2A and GluN2B subunits and a Fab domain. Colored bars are valid solutions while white bars correspond to false positives. Both locations of GluN1 are obtained from either copy despite conformational differences with the reference structure shown in yellow. The MaD score is significantly more effective to score the solutions related to the Fab domain. On the right, the first and best model obtained following the color code in the bar plots. **b**, Similar panel arrangement for the actin:tropomyosin complex (EMD-5751 at 8 Å, PDB 3J4K). The first and best model is shown. Descriptors from a single copy of the actin monomer led to correspondences at all 5 expected locations, enabling a fast assembly prediction. **c**, The VAT complex (EMD-3436 at 7 Å, PDB 5G4F) is similarly modelled from a single copy despite conformational differences (cRMSD) between protomers and missing densities. **d**, The ClpC-MecA complex (EMD-5609 at 10 Å, PDB 3J3U) was modelled without imposing symmetry. While a single and correct model was generated, the MecA subunits were too small for MaD to build reliable descriptors and find their location. **e**, The β-galactosidase in complex with variable regions of Fab domains (EMD-2548 at 13 Å, PDB 4CKD). At this resolution, the MaD score is more efficient than cross-correlation to discriminate false positives. MaD produced a single model with the expected stoichiometry, although variable antibody F_V_ domains could not be predicted.

In the next case, the asymmetric actin:tropomyosin complex at 7 Å, its five actin subunits are docked at their expected locations and ranked before two false positives (**Figure 2b**). The first of the two generated models exhibits a cross-correlation of 0.88 with the experimental density, a small improvement over the reference structure at 0.86. In the previous section, we mentioned that descriptor matching should naturally identify alternate locations of components of homomultimeric assemblies. The model shown here was obtained from a single subunit, whose descriptors were matched with those of the experimental map, enabling the prediction of the whole assembly from a single component (**Figure 2b**, right). The same is true even if protomers exhibit conformational differences. The homohexameric VAT complex with helical symmetry is fully modelled from a single monomer (used as example in **Figure 1** and **2**) despite root-mean square deviations of up to 3.38 Å between components and significant missing densities at some locations (**Figure 2c**). The unique model produced by our method shows a cross-correlation of 0.86.

At resolutions of 10 Å and lower, the densities of secondary structures elements tend to merge together. This is true in the case of the symmetrical MecA-ClpC complex at 10 Å (**Figure 2d**). Mad still generates reliable and discriminative descriptors, yielding a unique assembly model with the expected topology and cross-correlation of 0.92 to the reference structure, down from 0.94. However, details are not sufficient for the small MecA subunits, making it difficult to both extract stable anchors and build discriminative descriptors (**Figure 2d**, bottom). Regardless, this assembly was obtained without imposing the assembly’s C_6_ symmetry.

A similar result is obtained with β-galactosidase at 13 Å, a homotetramer with D_2_ symmetry with bound F_V_ antibody domains. The monomers are located correctly and the MaD score proved efficient at discriminating false positives, whereas the cross-correlation struggles to identify false positives (**Figure 2e**). However, the variable regions of antibodies could not be docked for similar reasons as the case of the MecA-ClpC complex.

Finally, our method was applied to a set of 21 assemblies simulated at a resolution of 10 Å and compared to recent similar methods^20,22,23^ (**Table 1**). MaD outperforms these other methods on this benchmark set, with the worst RMSD-Cα with the reference structure at 1.1 Å. Further, MaD produced a unique and correct model in all but one case (PDB 1Z5S). In the experimental cases described earlier, the first model always corresponded to the optimal one and at most 4 different models were generated. The fastest prediction took 7.78 seconds on a standard desktop workstation, while the slowest needed less than two minutes. As descriptor generation takes up about 60% of that time (**Table 1**), localizing components speeds up significantly past the first component since assembly descriptors are already computed. Further, descriptors are stored in a database, facilitating further modeling efforts should new component structures or cryo-EM data become available. Altogether, these results emphasize MaD’s ability to produce, score and rank models with unprecedented efficiency regardless of the target’s multimeric state, missing densities or small conformational differences.

**Table 1:** Performance of MaD on a set of 21 assemblies simulated at a resolution of 10 Å. Each row starts by the PDB access ID of the system, used as to simulate a map at a resolution of 10 Å. The rank is the position of the best model predicted by MaD over the total number of models produced. The number of subunits is then reported. The computation times in seconds for descriptor generation, descriptor matching and assembly building steps are shown, followed by the total prediction time. The last three columns report the root-mean-square deviation (RMSD) on Cα in Å between the reference PDB structure and the best predicted models by our method, IMP and γ-TEMPy, respectively. In the latter two cases, numbers have been taken from their respective articles^20,23^.

| System | Rank | #comp. | Computation time (sec) | | | | RMSD-Cα (Å) | | |
| --- | --- | --- | --- | --- | --- | --- | --- | --- | --- |
| | | | Gen. | Match | Build | Total | MaD | IMP | γ-TEMPy |
| 1cs4 | 1/1 | 3 | 12.49 | 4.87 | 0.25 | 17.61 | 0.50 | 2.60 | 2.80 |
| 1gpq | 1/1 | 4 | 6.05 | 2.56 | 0.59 | 9.20 | 1.10 | 1.70 | 0.60 |
| 1mda | 1/1 | 6 | 13.32 | 13.71 | 0.80 | 27.83 | 0.54 | 7.80 | 12.00 |
| 1suv | 1/1 | 6 | 31.99 | 24.81 | 1.35 | 58.15 | 0.26 | 5.20 | n/a |
| 1tyq | 1/1 | 7 | 28.41 | 17.46 | 0.56 | 46.43 | 0.45 | 19.10 | 16.30 |
| 1vcb | 1/1 | 3 | 6.14 | 1.49 | 0.14 | 7.78 | 0.67 | 1.80 | 4.80 |
| 1z5s | 1/2 | 4 | 8.76 | 2.96 | 0.24 | 11.95 | 0.50 | 8.70 | n/a |
| 2bbk | 1/1 | 4 | 11.51 | 5.30 | 0.50 | 17.31 | 0.84 | 2.10 | 11.30 |
| 2bo9 | 1/1 | 4 | 11.68 | 5.96 | 0.55 | 18.19 | 0.48 | 1.30 | 4.00 |
| 2dqj | 1/1 | 3 | 6.14 | 2.63 | 0.19 | 8.96 | 0.95 | 2.00 | 0.70 |
| 2gc7 | 1/1 | 4 | 12.69 | 5.23 | 0.27 | 18.19 | 0.59 | 1.30 | 10.90 |
| 2uzx | 1/1 | 4 | 20.37 | 7.31 | 0.86 | 28.53 | 0.28 | 1.10 | n/a |
| 2wvy | 1/1 | 3 | 21.41 | 14.98 | 0.61 | 37.00 | 0.23 | 0.90 | n/a |
| 2y7h | 1/1 | 3 | 21.86 | 7.75 | 0.70 | 30.31 | 0.25 | 1.70 | n/a |
| 3lu0 | 1/1 | 5 | 62.36 | 49.74 | 1.08 | 113.19 | 0.24 | 9.00 | n/a |
| 3nvq | 1/1 | 2 | 30.09 | 14.26 | 1.07 | 45.43 | 0.29 | 0.90 | n/a |
| 3pdu | 1/1 | 4 | 9.88 | 4.98 | 0.27 | 15.12 | 0.59 | 1.30 | n/a |
| 3puv | 1/1 | 5 | 28.19 | 28.58 | 0.82 | 57.59 | 0.43 | 22.70 | n/a |
| 3r5d | 1/1 | 3 | 9.26 | 2.18 | 0.24 | 11.67 | 0.82 | 1.40 | n/a |
| 3sfd | 1/1 | 4 | 16.91 | 6.49 | 0.43 | 23.83 | 0.51 | 1.40 | n/a |
| 3v6d | 1/1 | 2 | 19.75 | 7.76 | 0.29 | 27.80 | 0.29 | 1.60 | n/a |

### Effect of resolution on macromolecular descriptors

As resolution decreases, so does the amount of observable and useful details for building descriptors. At 6 Å, individual secondary structure elements should be distinguished, especially α-helices (**Figure 3a**). At 8 Å, β-sheets start to merge (**Figure 3b**). Above 10 Å, α-helices merge into more uniform densities but descriptors are still able to integrate shape information (**Figure 3c**). At 13 Å, only small domains may still be reliably described, severely limiting the amount of valid descriptor pairs. To ensure the capture of relevant shape information, the diameter of the patch may thus be increased (**Figure 3d**). To better quantify the effect of resolution on the performance of our method, we selected 100 assemblies with resolutions ranging from 5 to 15 Å originating either from single particle analysis or subtomogram averaging (**Supplementary Table 1**). The reference structure of each assembly was randomly displaced out of the experimental density and subsequently docked using our macromolecular descriptors. All but two assemblies reached a solution within 2 Å of the pre-fitted PDB (**Figure 3e**, top), which were at resolutions of 14 and 14.2 Å from single particle analysis. Importantly, valid solutions generally ranked first (**Figure 3e**, bottom) and exceptions usually correspond to symmetrical assemblies with several equally valid results. For all assemblies, we computed the number of valid descriptor pairs per residue in function of resolution (**Figure 3f**, top). As expected, this value decreases steadily with resolution. Furthermore, the pixel size tends to increase as resolution lowers. In practice, this means that lower resolutions targets are limited to appropriately sized subunits so that enough valid descriptors may be generated and reliably matched. During the global matching step (**Figure 1e**), four descriptor pairs agreeing with a localization is imposed to validate a candidate and limit false positives. Taking this observation into account, the minimum molecular weight of components to be docked as a function of resolution may be estimated from the latter distribution (**Figure 3f**, bottom). This results in the ability to locate subunits the size of ubiquitin (ca. 8 kDa) at resolution close to 5 Å, while 60 kDa and bigger structures would be required for successful predictions at resolutions nearing 15 Å.

**Figure 3:**
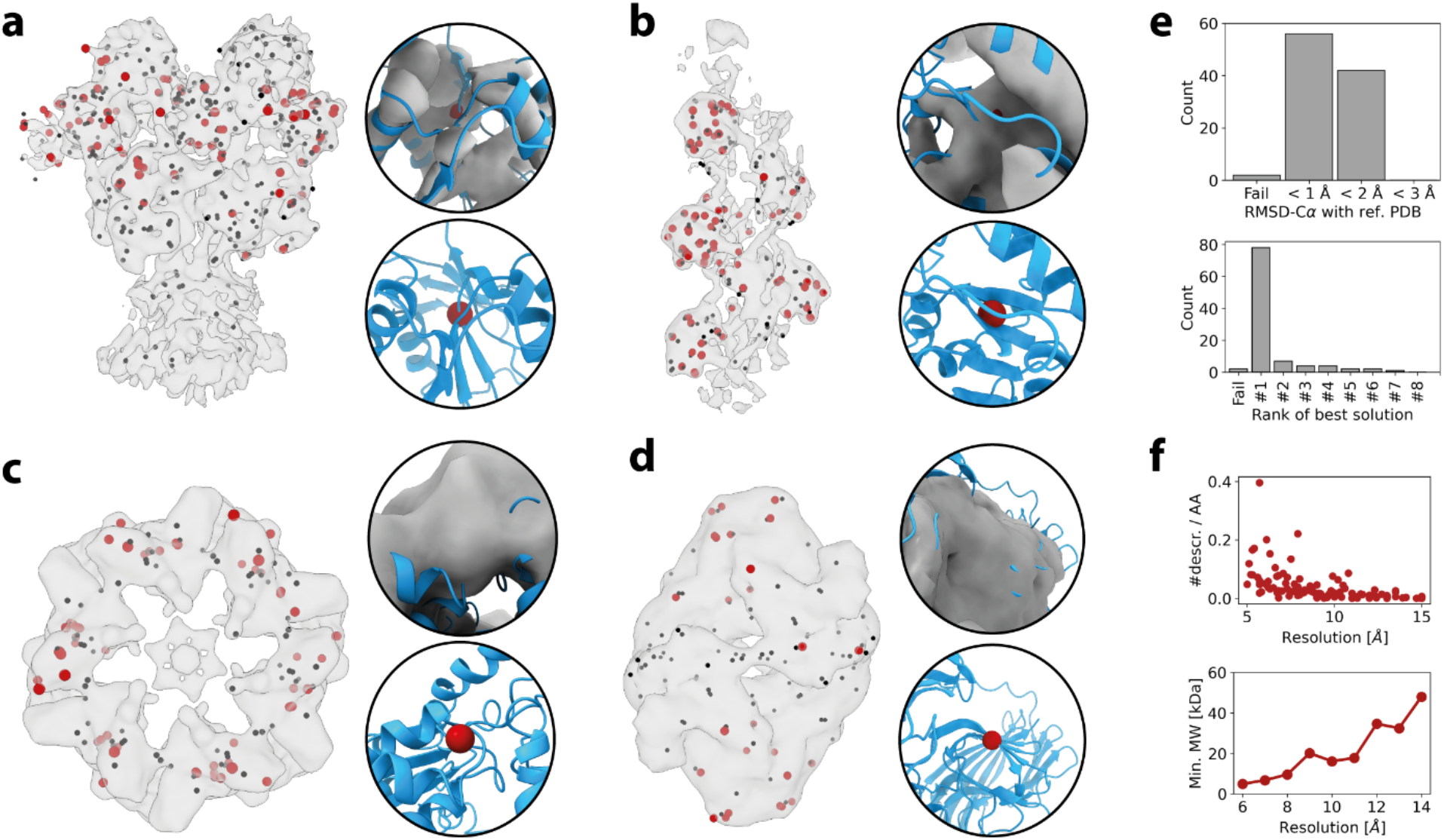
Effect of resolution on descriptor performance. In panels a-d are shown anchors that lead to valid descriptor pairings (red) and those that did not produce valid descriptor pairings but counted as inlier during the matching procedure (black). Structural regions corresponding to descriptor pairs that lead to valid localizations are shown on the right side of panels a-d. **a**, NMDA receptor at 6 Å. **b**, actin:tropomyosin at 8 Å. **c**, MecA-ClpC complex at 10 Å. **d**, β-galactosidase at 13 Å. **e**, results achieved by docking 100 structures in cryo-EM maps obtained by either single particle analysis or subtomogram averaging at resolutions between 5 and 15 Å. RMSD-Cα with the reference structure and the rank of the best model are shown. **f**, the number of valid descriptors per amino acid (AA) residue in function of resolution for the assemblies of panel **e** that were successfully docked. On the bottom is shown the minimum molecular weight of structures in function of resolution estimated from the above panel.

### Including structural ensembles for a dynamic modeling of GroEL

Macromolecular assemblies are inherently flexible and explore complex conformational landscapes in order to fulfil their task in the cell. Furthermore, components may exhibit different behaviours in isolation or in their oligomerised state, conformational changes occurring upon assembly notwithstanding. As such, hybrid modeling strategies should account for the fact that the structures being integrated are not necessarily in a compatible conformational state. Consequently, it is crucial that modeling methods be able to consider structural ensembles and to detect compatible structures within it. We previously showed that valid ensembles are obtainable through molecular dynamics and may be explored efficiently by an adequate optimizer^12–14^. While the interconnectivity between timesteps may be favourable for the convergence of continuous optimizers, MaD is, in essence, a discrete docking software. Therefore, the inherent redundancy of molecular simulations is detrimental to its performance and a pre-processing step is necessary to efficiently include dynamic information. We recently published a clustering algorithm, CLoNe, able to tackle large and complex conformational ensembles from various contexts^40^. In particular, its density estimation and processing enable the reduction of trajectories into an ensemble of cluster centers, which may be viewed as highly populated and unique states in accordance with the sampled conformational landscape. Combined with macromolecular descriptors, the resulting pipeline, termed MaD-CLoNe, is one that is able to consider large ensembles without wasting extensive time on redundant states or those incompatible with the target cryo-EM data. As previously mentioned, cryo-EM descriptors are computed only once for all components or all structures of an ensemble, emphasizing the ability of MaD to consider large assemblies and structural ensembles with a minimal increase in computation cost and complexity.

To demonstrate this, let us consider the tetradecamer of GroEL (EMD-5338 at 6.1 Å) (**Figure 4a**), whose subunits are each made up of an ATP-binding equatorial domain, catalytic intermediate domain and a substrate, GroES binding domain^41^. We applied our clustering method (i.e., CLoNe) to a 200 ns classical molecular dynamics simulation of a GroEL monomer after projecting it onto the first two principal components and identified 7 relevant cluster centers (C1 to C7 in **Figure 4b**). We then docked the resulting ensemble of centers (**Figure 4c**) into the density map using MaD. By averaging the scores of the solutions obtained by each independent structure, the best candidate is naturally made evident (**Figure 4d**). Of note, the MaD score correctly estimates the conformational compatibility with the target cryo-EM data, where C5 and C1 are the closest and farthest to the reference structure, respectively (**Figure 4de**). Our method identifies exactly 14 solutions for C5, which amount exactly to the correct topology with a cross-correlation of 0.79 with the target density, while valid descriptors are identified over the whole structure (**Figure 4f**). In contrast, 30 solutions are found for C1. The best assembly still respects the expected tetradecameric topology with a cross-correlation of 0.57 (**Figure 4g**), with valid descriptors located exclusively to the equatorial domain. While two of the remaining solutions are false positives, the other 14 correspond to another valid assembly, albeit with significant internal clashes (**Figure 4h**), with valid descriptors located in the apical and intermediate domains instead. As a result, the descriptor scoring enables, at a glance, the identification of the best candidates from a structural ensemble. Furthermore, this demonstrates the robustness of our method towards strong conformational differences, in which case it also provides insight into the different ways of fitting components within the target density, potentially guiding further modeling steps.

**Figure 4:**
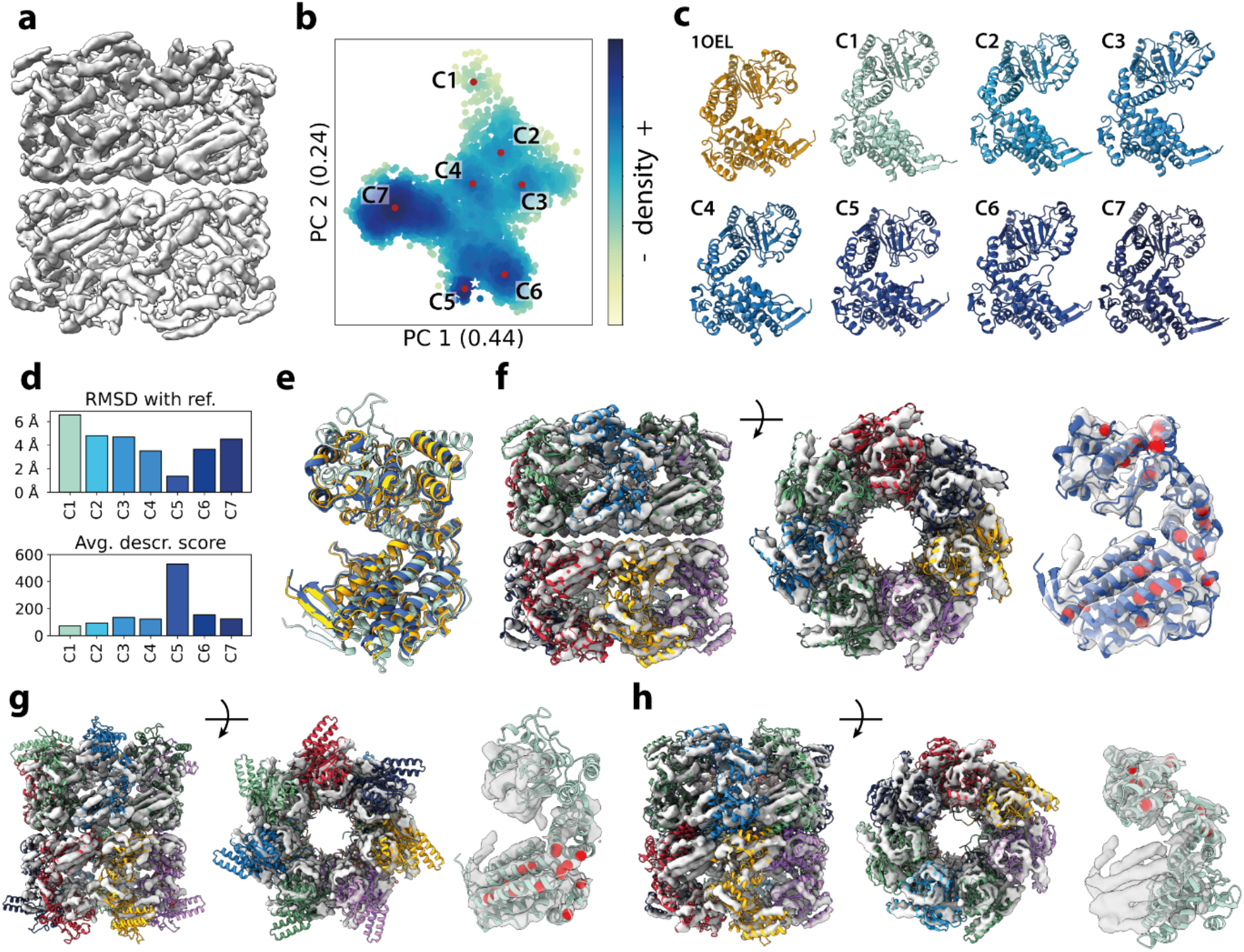
Dynamic integrative modeling of GroEL. **a**, the target experimental density map EMD-5338 at 6.1 Å. **b**, projection of a 200 ns classical molecular dynamics simulation with over 5000 structures onto two conformational coordinates identified by principal component analysis, whose eigenvalue are of 0.44 and 0.24, respectively. Cluster centers are shown as red dots and labelled from C1 to C7, with the reference structure shown as a white star. **c**, the reference structure used an initial state for the simulation and the identified cluster centers. **d**, the RMSD-Cα between the cluster centers and the reference structure (top) and the average MaD score obtained by each cluster center. **e**, the reference structure (orange) superimposed with the best scoring conformation, C5 (dark blue) and the worst one, C1 (light blue). **f**, the assembly obtained from C5 with the highest score and highest compatibility with the cryo-EM data. On the right, a close-up view of a with the valid descriptors shown as red spheres. **g**, best assembly obtained from the lowest scoring conformation, C1, with a close-up view showing valid descriptors. **h**, the alternate assembly obtainable from C1 with a different alignment to the density, with valid descriptors shown on the right side of the panel.

## Discussion

This work introduces MaD, a hybrid modeling approach inspired by computer vision concepts able to predict assemblies on the sole basis of non-atomic cryo-EM maps and high-resolution structures of the components, or their high-confidence models which are now accessible thanks to the recent progresses in protein structure prediction.^5,6^ The appeal of our work lies in its easy identification of solutions, high robustness to conformational differences between input structures and its ability to process and scale to structural ensembles efficiently. Compared to methods relying on stochastic optimizers, our method in general does not require several independent runs to guarantee optimal results. Moreover, the descriptor-based scoring is valid even at supra-nanometer resolutions where cross-correlation loses its reliability. Importantly, our score reports on the conformational compatibility between input structures, enabling the easy identification of the best candidates for assembly prediction. This is a distinct advantage over traditional scoring functions, which include different terms often incompatible with structural mismatch, such as protrusions from the target experimental density or shape complementarity between components. When compatible structures are not available, our method might provide insight into different docking positions, helping the modeling process rather than bringing it to a halt.

Despite the tremendous growth of the EMDB, most integrative methods still report results on simulated data. Simulated densities however pale in comparison of experimental data and the different methods or factors used in their generation may vary. Importantly, no conformational mismatch between input structures is present, which may lead to unrealistic results. Here, we only utilized simulated assemblies for comparison purposes and suggest that experimental data should be used exclusively for further method development. We tested our method up to resolutions of 15 Å. Structural details at even lower resolutions may not be sufficient for reliable descriptors to be generated and as such, we expect other methods to be superior to ours in such situations. We however expect that the relevance of these resolutions will continue to decrease following the resolution revolution and as further progress is achieved in both single particle analysis and subtomogram averaging. While the average resolution of single-particle cryo-EM has been stagnating around 5 Å in recent years, the one of subtomogram averaging reached 20 Å in 2021, with further potential from hybrid reconstruction methods^2^. Should it continue to evolve positively, we expect MaD to be a prime candidate for modeling large macromolecular assemblies in their native cellular environments.

Further, descriptors embody a concept that is applicable at multiple scales as long as data is represented as evenly-spaced grids. In practice and possibly in conjunction with future progress in super-resolution microscopy, proteomics and de novo prediction of molecular complexes, we may envision promising applications towards the painting of high-resolution images of whole cellular environments by docking assemblies in tomograms rather than components in assembly maps. On the other side of the molecular scale, small-molecule docking software may benefit from descriptors as well, especially as key and recent approaches adopt a fragment-based methodology. In light of these results, MaD constitutes a solution to current issues in integrative modeling and exhibits potential to remain relevant for years to come thanks to the adaptability of its core concepts.

## Methods

### Anchor detection

High-resolution component structures are projected onto a grid of identical spacing as the target experimental cryo-EM data. The resulting grid is convolved with a Gaussian kernel with a σ factor of 0.225, identical to the default value in Chimera’s *molmap*. Voxel values are then normalized between 0 and 1. In 2D, many descriptors are scale-invariant because objects may appear at different sizes between images. To this end, a Gaussian pyramid is commonly built to extract features^35,42^. Pyramid levels, otherwise referred to as octaves, include a specific image size with different levels of blurring. An octave is defined by the size of the image, where subsequent levels are half the size of the previous one. As biological structures are encoded in real-world metrics, a unique scale is needed. In practice, we generate two single-scaled octaves, with the first corresponding to the original grid and the second to a copy supersampled by a factor 2 to retrieve some higher frequency features lost to the blurring. After convolution with a Laplacian of Gaussian, anchors are detected as voxels with higher values than their 26 neighbours. Anchors with values below 5e^-2^ are discarded due to poor contrast. Saddle points are filtered by checking the eigenvalues of the Hessian matrix. A strong curvature is often ensured with a threshold on the ratio of the largest and smallest eigenvalues, but this procedure did not improve detection in our case. Finally, anchor coordinates are refined to subvoxel accuracy using a Taylor expansion similarly to SIFT^35^:

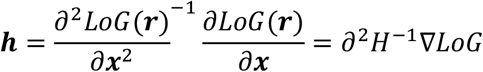

where ***h*** is the offset to apply to the anchor’s coordinate. If the offset is larger than 0.5, the anchor is displaced to the nearest voxel and the procedure is repeated. If no convergence is achieved, the anchor is discarded for being unstable.

We aimed for a detection scheme invariant to resolution. To this end, we measured repeatability, defined as the fraction of anchors from an atomic structure having a corresponding anchor in its experimental cryo-EM map. Such a correspondence was accepted if the anchors were at most one voxel apart, diagonal included. We tested our detection scheme on 6 systems at resolutions of 5 to 10 Å, using either experimental or simulated cryo-EM data (**Supplementary Figure 1**). Using a supersampled octave presmoothed with Gaussian of scale σ=1 yielded higher correspondences and repeatability than using the original grid alone or both octaves separately, result that is in line with 2D procedures^35^. As expected, experimental data performs worse than simulated data due to noise and systematic errors, but sufficient anchors are detected in all cases presented. By default, σ=2 was chosen and used for all results presented despite the maximum of correspondences generally found at σ=1.5. This choice was made based on the observation that a subset of correspondences is sufficient for subsequent steps and thus lower values only lead to longer computing times.

### Orientation assignment

Around each detected anchor, a patch is extracted as to include sufficient information from secondary structure elements. This value ultimately depends on the resolution and pixel size the target cryo-EM data, which tend to be between 1 and 1.5 Å for subnanometer resolution maps (see **Supplementary Table 1**). By default, a patch of about 25 Å in diameter is used. As patches are not rotation invariant at this point, corners are disregarded to prevent bias. In contrast to 2D descriptors, at least two angles are necessary to provide rotation invariance to an anchor in 3D. While orientation histograms are easily built in ℝ^2^ to sort gradient vectors according to their direction and magnitude, the situation is more complex for ℝ^3^. Previous 3D descriptors divided the unit sphere into evenly spaced angular intervals across the polar and azimuthal angle^32,43^, but this produces an unequal division of the sphere. The only exact solutions to dividing the sphere into equal areas correspond to the platonic solids, of which the icosahedron is used in combination with an eigen-decomposition of the structure tensor in a generalisation of descriptors to ℝ^*n*31^. However, the icosahedron only has 20 faces, while SIFT descriptors utilize 36 bins already in ℝ^2^. As such, this approach may lack accuracy in deriving rotation matrices for biological structures at low resolution. Instead, the zonal equal sphere partition scheme (EQSP)^39^ is particularly well-suited for building orientation histograms. EQSP spheres have an arbitrary number of zones of approximately equal area and shape, each defined by a center and the angular range it covers, with poles represented by circular caps (**Figure 1b**, second column and **Supplementary Figure 2d**). As such, they represent an ideal medium to sort the gradient vectors according to their direction using range-based assignation. The zone with the highest vector count yields the first dominant orientation. Orientation assignment is usually done by weighing vectors according to their magnitudes. However, we found that simply counting vectors is faster and integer-based, and the performance loss compared to weighing their contribution with their magnitude was negligible. The patch is rotated along the polar angle φ so that the dominant zone is located on the upper cap of the EQSP sphere, yielding the rotation matrices *R*_*c*Φ_ and *R*_*a Φ*_, for the component and assembly anchor, respectively. The axis of rotation is obtained by the cross-product between the gradient vector and the Z axis. The second orientation is obtained by repeating the process but excluding both caps and rotating about the Z axis at an angle θ this time so that the bin is the first of its layer, yielding the matrices *R*_*c*θ_ and *R*_*a*θ_. For both orientations, any ambiguous bins with a vector count that is at least 80% of the maximum is considered by duplicating the anchor accordingly. Rotating a component *c* to its preferred orientation in the context of the experimental cryo-EM map of assembly *a* can then be done using the following matrix:

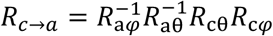

Combined with a translation vector *T*_*c*→*a*_ between anchor coordinates, this rotation matrix enables the docking of a component in the target density provided that their anchors match in location and orientation.

To benchmark this step, we selected a number of subunits of different shapes and sizes (**Supplementary Figure 2a**). The following parameters were considered: the size of the extracted patch in voxel, the scale of a Gaussian window centered on the anchor to weigh voxels based on their distance to the anchor and the number of zones on the EQSP sphere (**Supplementary Figure 2bcd**). The number of zones were chosen so that each sphere exhibits an additional layer of zones. Patch size was limited to 17 voxels maximum to not include excessive blank space around smaller subunits. The benchmarking procedure was focused on repeatable anchors only and aimed to maximize repeatable anchors with matching orientation. This can be measured by applying the transformations *T*_*c*→*a*_ and *R*_*c*→*a*_ from each anchor pair and measuring the RMSD-Cα with the fitted component. In practice, if the anchor pair is located on one side of an elongated structure, this measure may be large despite being valid. Thus, we instead enforce an angular displacement of at most 30° between the docked and reference structures, as the translation is valid by definition. Optimally, this procedure should be independent of the original placement and orientation of the component in space. To account for this, the fitted component was translated away from the density and rotated at regular intervals of 15° from 0° to 360° around all three Euler axes simultaneously prior to fitting.

Benchmark results show that a patch of 17 voxels in diameter, an EQSP sphere with 112 bins and no Gaussian window yield the highest amount of repeatable anchors with matching orientations (**Supplementary Figure 3a**). Both the minimum RMSD-Cα with the reference structure and percentage of valid anchor pairs is stable over rotation (**Supplementary Figure 3b** and **c**, respectively). Only the ClpC subunit of PDB 3j3u showed larger fluctuations in RMSD-Cα due to its elongated shape and lower resolution of the target density map at 10 Å. A similar procedure was repeated for whole structures on a set of 60 cryo-EM maps between 5 and 10 Å rather than individual components (**Supplementary Figure 4** and **Supplementary Table 1**). This highlighted that a larger patch diameter may be used for a slightly higher efficiency should the structure be large enough. The other parameters are in line with the results of singular components.

### Descriptor generation and local matching

The identification of valid anchor pairs is facilitated by providing them with a robust representation in the form of unidimensional vectors. To this end, the patch around an anchor is resampled after it has been assigned an orientation. Since patches are rotation invariant at this point, corners are considered as they are no longer a source of bias but may serve to discriminate valid pairs. The resampling is done through trilinear interpolation and performed so that values are evaluated at positions in between existing voxels, allowing the patch to be equally subdivided in a number of subvolumes (**Supplementary Figure 5a**). The subdivision of the patch enforces that relevant information is not only present in the patch, but at a specific location around the anchor, enhancing discriminative power. Further, this allows the information to shift within a given subvolume without impacting the final descriptor, providing robustness to localization shifts due to noise and conformational differences. Within each subvolume, gradient vectors are computed at all positions and sorted according to their direction on an EQSP sphere, which contains fewer zones than during orientation assignment (**Supplementary Figure 5b**). This allows for small shifts in gradient orientation while accounting for the fact that fewer vectors are present in a subvolume. At this point, the patch is encoded by a list of EQSP spheres summarizing the information from their respective subvolumes. The corresponding histograms are all flattened in a single vector and normalized, yielding the final descriptor. A single descriptor thus has *nr* values, where *n* is the number of subvolumes and *r* the number of zones on the EQSP sphere.

In 2D, a common filtering procedure of descriptor pairs involve the nearest-neighbor distance ratio (NNDR)^35,44^. The aim is to find the two closest neighbours of a descriptor from a first image within the descriptors if another image. If the match with the first neighbour is genuine, the ratio of the two should be low. If not, that ratio should be closer to 1 as both neighbors are far from the query descriptor. Depending on the situation, neighbors further than the second one may be considered. However, biological structures are repetitive entities, either within a component depending on secondary structure repetition or in the case of homomultimers, making this approach poorly applicable without tedious fine-tuning. Instead, we treat the dot product between normalized descriptors as a probability of that pair being a genuine match. This allows the setting of a confidence threshold above which a descriptor pair is accepted as candidate.

To ensure that our procedure rank valid descriptor pairs first, we looked for parameters maximizing the average area under the receiver operating characteristic (AUROC) over the same angular intervals and range as in orientation assignment. In practice, relevant parameters include the number of subvolumes to extract from the patch around the anchor and the number of zones of the EQSP sphere assigned to each subvolume (**Supplementary Figure 5ab**). As expected, a higher number of subvolumes lead to more discriminative descriptors (**Supplementary Figure 5c**, left) and the 4×4×4 subdivision is chosen by default. The size of the spherical histogram however is not significant above 8 bins (**Supplementary Figure 5c**, right). For a patch diameter of 16 voxels after interpolation, a given subvolume has 64 gradient vectors. Thus, we enforced 16 zones on each EQSP sphere during descriptor generation to better allow for small shifts in gradient orientation without significantly impacting performance and avoid overfitting. Importantly and similarly to orientation assignment, the AUROC is robust to the initial position of the component (**Supplementary Figure 5d**).

### Global matching

Descriptor matching is only a local measure of similarity which does not translate to whole structures on its own. SIFT employs a Hough transform^35^, so that selected descriptor pairs may vote for specific locations in an image. In practice, random sample consensus (RANSAC)^45^ or even progressive sample consensus (PROSAC)^46^ may be used. However, they work best for a relatively high number of inliers, which is not always guaranteed especially if conformational differences are present. Contrary to many descriptor applications, another important aspect of hybrid modeling is that more than one solution is often expected as specific components may be present in multiple copies in the target assembly. To that end, we combine concepts from both the Hough transform and PROSAC^46^, enabling identified descriptor pairs to vote for several locations at once while processing a fraction of the pairs. PROSAC^46^ orders samples according to a quality function *q*, defined here as the repeatability between the component’s anchors and those of the assembly after applying the *T*_*c*→*a*_ and *R*_*c*→*a*_ associated to that pair. Each pair of the ordered list then votes for a component localization within the target assembly, resulting in a fast hierarchical clustering step proceeding as follows. The first descriptor pair with highest repeatability is defined as a cluster seed and its associated *T*_*c*→*a*_ and *R*_*c*→*a*_ are applied to the component’s anchors. The same is done with the next pair. It is either assigned to the first cluster if the anchor cloud is located within an RMSD 10 Å of the first seed or a new cluster is created and this pair becomes a new seed instead. Subsequent pairs are processed similarly, either enriching a previous cluster or seeding a new one. The benefit of ordering pairs according to repeatability is two-fold. One, the first seed of a cluster may be safely assumed to be the best of the cluster. Two, the pairs leading to expected solutions should be the first to be processed and the clustering can be truncated, speeding up the procedure and limiting the inclusion of false positives at the same time. By default, the best 60 descriptor pairs are considered per component, value that is then multiplied by the number of copies of that component in the assembly. This was determined empirically and used for most of the data presented. It may be modified as needed, for example depending on the resolution of the target assembly. Namely, it was increased to 100 and 120 for the MecA-ClpC complex at a resolution 10 Å and β-galactosidase at 13 Å, respectively, as to obtain all expected solutions.

The size of clusters is interpreted as the weight of a particular solution. A minimum of 4 descriptor pairs is enforced to filter out most if not all false positives. True positives are often enriched not only by more unique anchors, but also by descriptors generated from ambiguous orientations. Conversely, false positives usually exhibit few agreeing pairs that may not make the list of processed pairs. Finally, the extracted solutions are refined using a rigid optimization alternating small rotational and translational steps similar to the procedure Fit-In-Map of Chimera^28^. After the refinement procedure, it is possible that different solutions became identical if the RMSD between their anchor cloud was just above the threshold during the clustering step. A simple pass through the solutions is performed to group identical solutions together, in which case their score is combined. The repeatability is also updated, as after refinement more correspondences are usually present. The final MaD score *S* for solution *i* is the product of different measures:

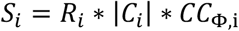

where *R*_*i*_ is the repeatability or percentage of corresponding anchors, |*C*_*i*_| the size of the cluster of agreeing descriptor pairs, and *CC*_Φ,i_ the cross-correlation with the target experimental cryo-EM data. As such, this score measures not only the global agreement between densities and the correspondence between anchors but also their surrounding patch, yielding a robust scoring incorporating information at multiple levels.

### Assembly building

As shown extensively in the results section, our method produces few false positives for individual components. The process of building assemblies from a list of solutions of components is thus straightforward in general and can be limited to a simple combinatorial search. In the general case, two steps are performed. First, a table of pairwise overlap between the solutions is computed. Overlap is defined as the number of non-zero voxels at the same point in space between two density grids divided by the amount of non-zero voxels. To this end, components are converted to simulated density grids at a resolution of 5 Å and voxel spacing of 2 Å. A minimum contour level of 0.2 is imposed as well after fixing the values between 0 and 1 in the grid. Then, for every possible assembly with the expected stoichiometry, the sum of overlap is computed. The selected models are ranked according to this value and any assemblies with more than 10% of overlap is discarded. In the case of partly homomultimeric assemblies, an intermediate step is added to reduce combinatorial complexity. Sub-complexes are built from the solutions of repeated components alone. Then, the sub-complexes are treated as individual components and the previous method is applied to generate the final models.

## Code availability

Macromolecular Descriptors (MaD) has been coded in Python3 as stand-alone. Code, basic instructions and required packages are available from the website of the Laboratory for Biomolecular Modeling (https://www.epfl.ch/labs/lbm/resources/) or on Github (https://github.com/LBM-EPFL/MaD).

## Supplementary information

**Supplementary Figure 1:**
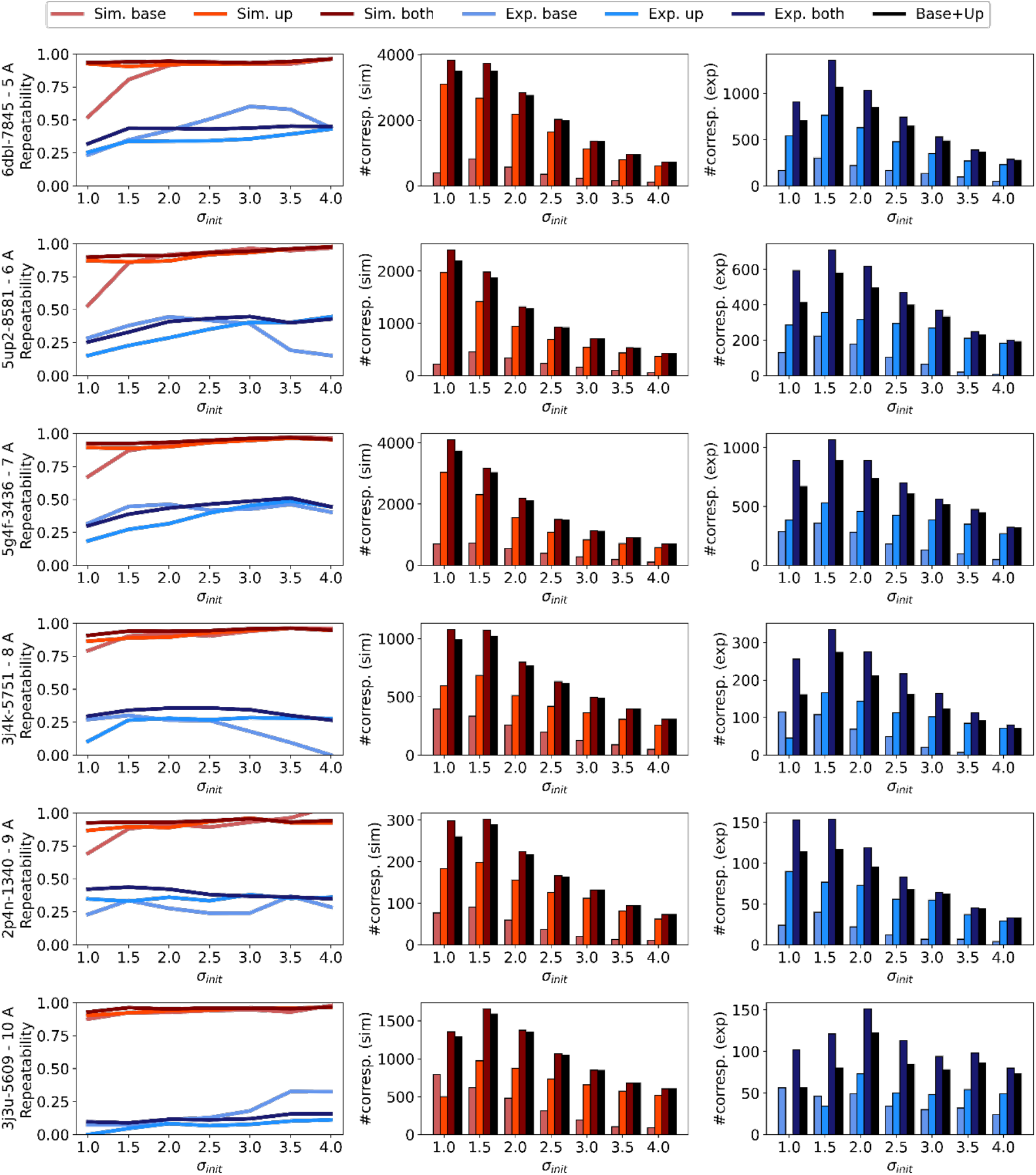
Anchor detection with Laplacian of Gaussian. Each row corresponds to a system. The PDB and EMD access IDs are shown, starting with PDB 6dbl and EMD 7845 at a resolution 5 Å and ending with PDB 3j3u and EMD 5609 at 10 Å. The first column shows anchor repeatability as defined in the Methods section, for the base, supersampled and both octaves simultaneously at different values of σ_*init*_. Both octaves have a single scale, filtered with a Laplacian of Gaussian of scale σ_*init*_ with the supersampled one pre-filtered with a Gaussian with scale σ = 1. Results obtained with the simulated map are shown in shades of red, and those obtained with the experimental map in shades of blue. In the second and third columns are shown the number of correspondences with simulated and experimental maps, respectively. The black bars are the sum of the correspondences obtained individually in the base and supersampled octaves.

**Supplementary Figure 2:**
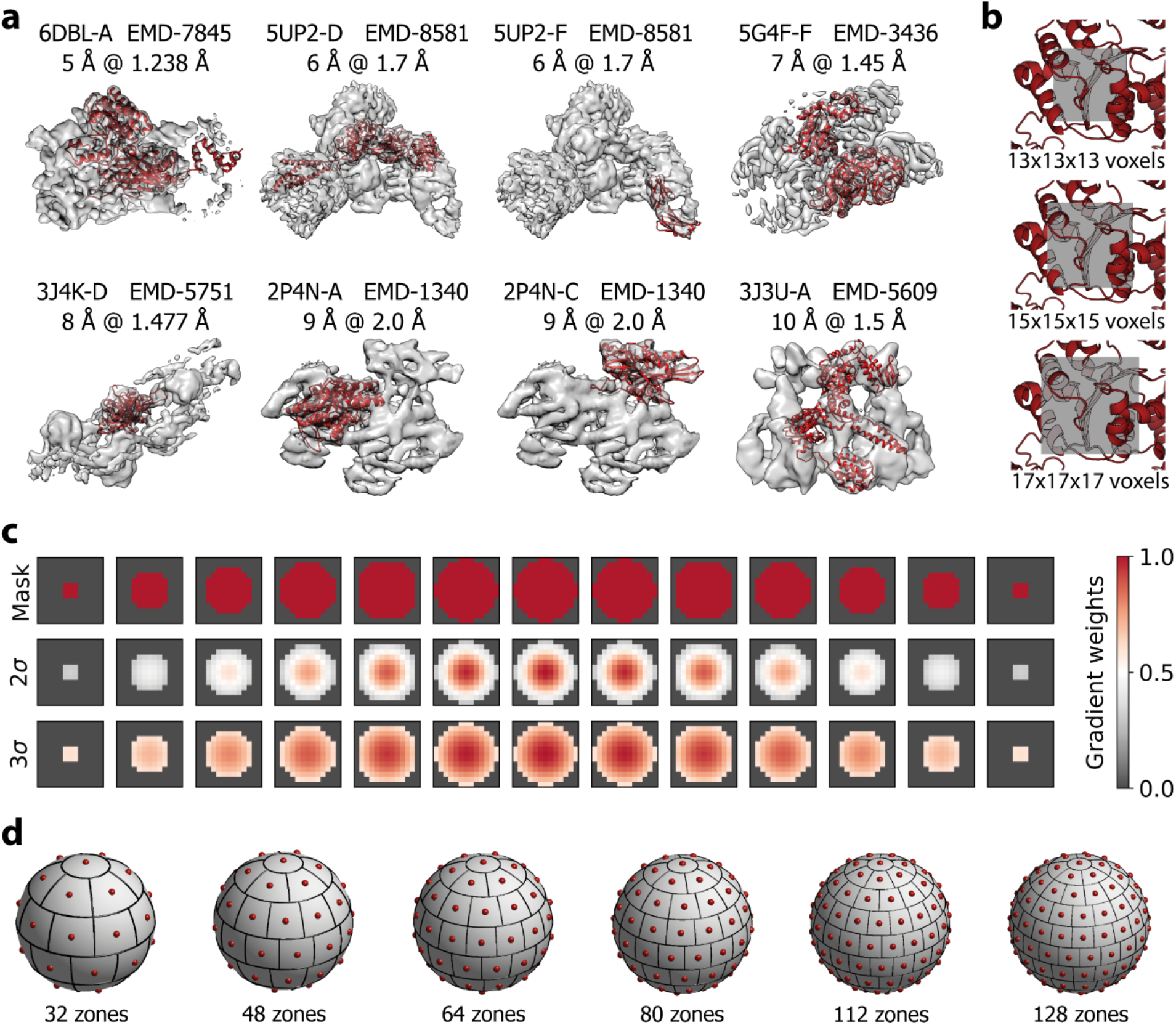
Benchmark parameters and structures for orientation assignment. **a**, the subunits and respective assemblies used in the benchmark, showing the PDB access code and the selected chain as well as the EMD number, resolution and voxel spacing of the corresponding experimental map. **b**, different sizes of the patch extracted around an anchor. Values shown refer to the diameter, including the planes crossing at the location of the anchor. A commonly found voxel spacing of 1.5 Å was used to render the patch. **c**, the mask used to extract the spherical patch from the cubic region is shown. Each image is a slice from top to bottom, where the considered voxels are colored in red. Each contribution may be weighted by a Gaussian function centered on the anchor, so that contribution for points far from the center are diminished. The σ of the Gaussian corresponds to the one used in the anchor detection step. **d**, the different EQSP sphere sizes considered.

**Supplementary Figure 3:**
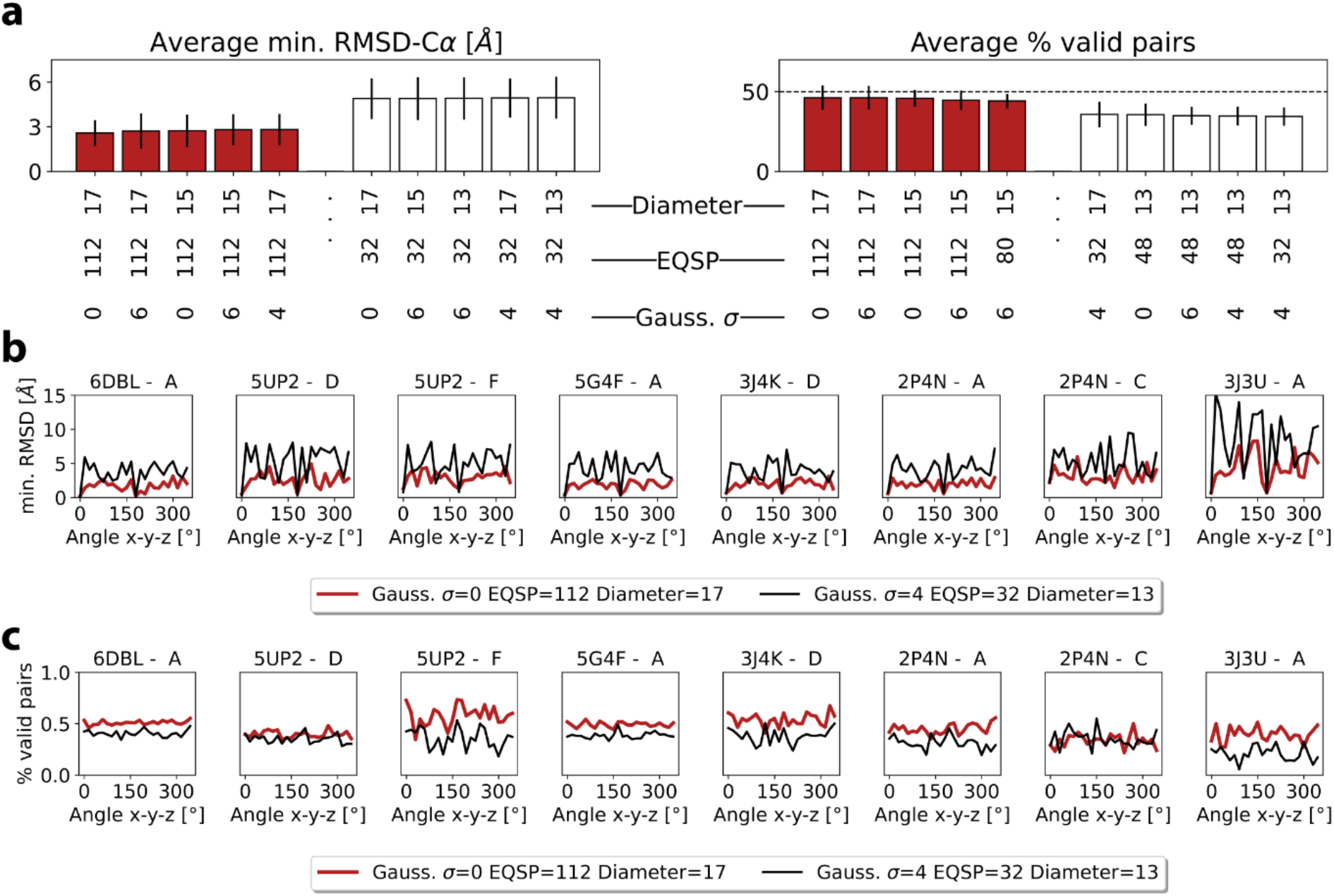
Benchmark results for orientation assignment. **a**, rankings of parameter sets, including the patch diameter in voxels, the number of EQSP zones and the relative σ of the Gaussian window, expressed as a multiple of the value used in the detection step (σ = 2). The best parameter sets are shown in red and the worst ones in white. **b-c**, The minimum RMSD-Cα with reference and fraction of valid anchor pairs are shown for each angles between 0 and 360° with a step of 15°. The best parameter set is shown in red, and the worst in black.

**Supplementary Figure 4:**
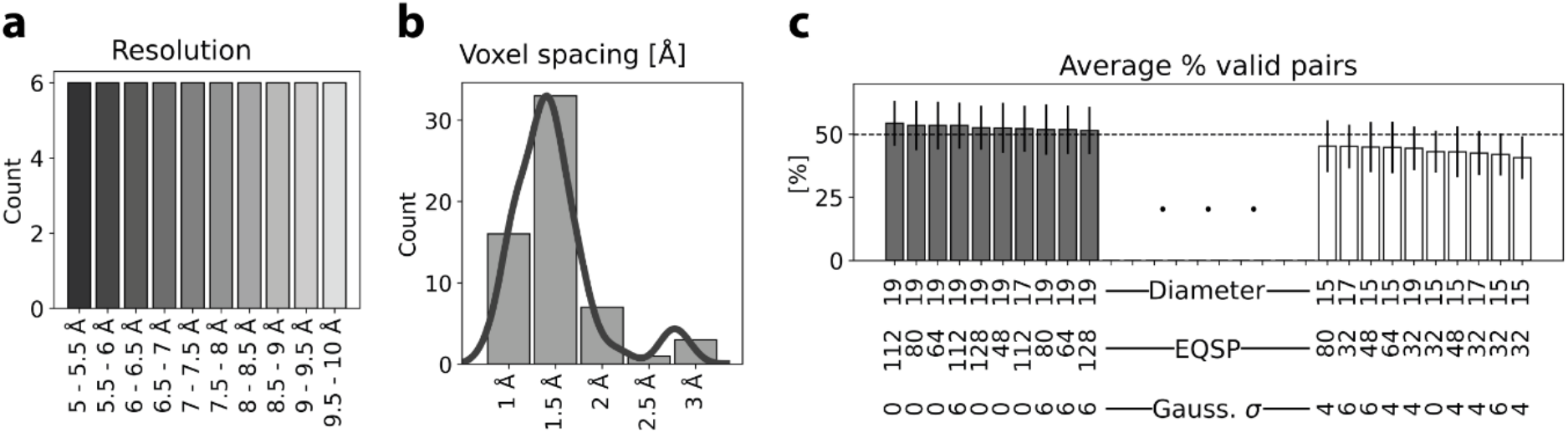
Benchmark results for orientation assignment on whole assemblies. **a**, the selected 60 assemblies sorted by resolution range (see Supplementary Table 1). **b**, the distribution of voxel spacing. **c**, ranking of parameter sets, including the patch diameter in voxels, the number of EQSP zones and the relative σ of the Gaussian window, expressed as a multiple of the value used in the detection step (σ = 2). The best parameter sets are shown in grey and the worst ones in white.

**Supplementary Figure 5:**
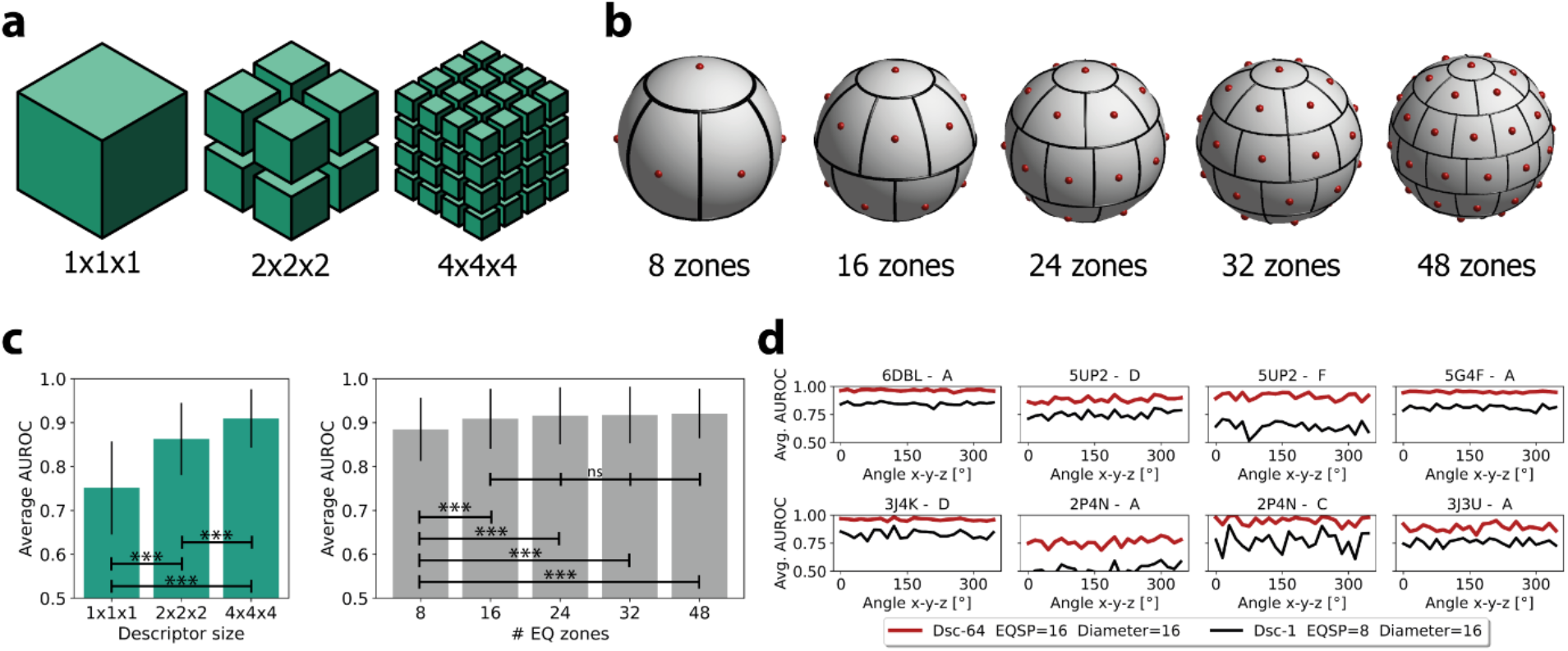
Benchmark for descriptor generation. **a**, the density patch around an oriented anchor may be kept as is, or divided into 8 (2×2×2) or 64 (4×4×4) subregions after an interpolation step. **b**, the different number of zones of the EQSP sphere. Smaller sizes than in orientation assignment are tested due to the lower number of voxels per subregion. **c**, On the left, the average area under the receiver operating characteristics curve (AUROC) is shown for each subdivision scheme. On the right, the average AUROC computed in the same way but for the number of zones on each EQSP sphere. p-values below 0.001 obtained by T-tests are indicated by ***, while non-statistically significant differences are denoted by ns. **d**, The AUROC accomplished per subunit for a 4×4×4 subdivision scheme with 16 EQSP zones compared to a descriptor without subdivision and an EQSP sphere with 8 zones, corresponding to the worst performance achieved.

**Supplementary Table 1:**
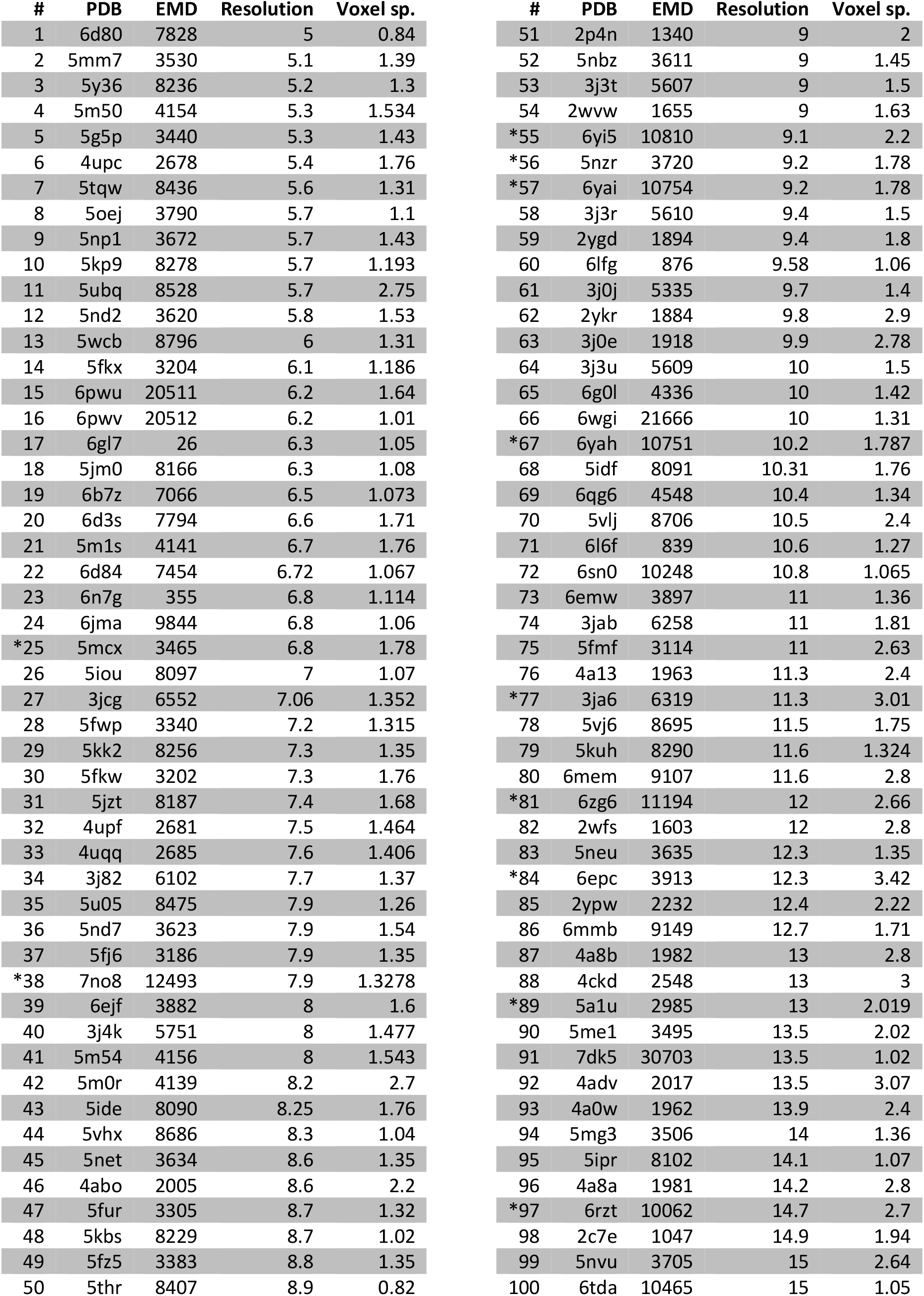
List of assemblies used in Figure 3 and Supplementary Figure 4. PDB and EMD access codes are listed. Resolution and voxel spacing are reported in Å. Systems have been reconstructed using subtomogram averaging are marked with *. Otherwise, single particle analysis was used. For Supplementary Figure 2, the first 60 assemblies using single particle analysis up to PDB 6g0l were used.

